# SARS-CoV-2 triggers DNA damage response in Vero E6 cells

**DOI:** 10.1101/2021.09.08.459535

**Authors:** Joshua Victor, Jamie Deutsch, Annalis Whitaker, Erica N. Lamkin, Anthony March, Pei Zhou, Jason W. Botten, Nimrat Chatterjee

**Affiliations:** Department of Microbiology and Molecular Genetics, Larner College of Medicine, University of Vermont, Burlington, VT 05405; Division of Immunobiology, Department of Medicine, Larner College of Medicine, University of Vermont, Burlington, VT 05405; Department of Biochemistry, Duke University School of Medicine, Durham, NC 27710; University of Vermont Cancer Center, University of Vermont, Burlington, VT 05405

**Keywords:** SARS-CoV-2, DNA damage response, genome instability, telomeres

## Abstract

The novel severe acute respiratory syndrome coronavirus 2 (SARS-CoV-2) virus responsible for the current COVID-19 pandemic and has now infected more than 200 million people with more than 4 million deaths globally. Recent data suggest that symptoms and general malaise may continue long after the infection has ended in recovered patients, suggesting that SARS-CoV-2 infection has profound consequences in the host cells. Here we report that SARS-CoV-2 infection can trigger a DNA damage response (DDR) in African green monkey kidney cells (Vero E6). We observed a transcriptional upregulation of the Ataxia telangiectasia and Rad3 related protein (ATR) in infected cells. In addition, we observed enhanced phosphorylation of CHK1, a downstream effector of the ATR DNA damage response, as well as H2AX. Strikingly, SARS-CoV-2 infection lowered the expression of TRF2 shelterin-protein complex, and reduced telomere lengths in infected Vero E6 cells. Thus, our observations suggest SARS-CoV-2 may have pathological consequences to host cells beyond evoking an immunopathogenic immune response.

## Introduction

SARS-CoV-2 is a novel coronavirus, and the causative agent of the COVID-19 pandemic. SARS-CoV-2 is a zoonotic positive-sense RNA virus from the *Coronaviridae* family of viruses. Since the inception of the pandemic more than 4 million deaths have occurred and some recovered patients have continued to report debilitating, and sometimes new symptoms long after the infection has ended. This condition with lingering symptoms is often referred to as “Long COVID”. Although mechanisms are unclear, hypotheses include viral-induced tissue and organ injury, in which SARS-CoV-2 infection alters host cell physiology, and cellular functions might be permanently altered. Hence there is an urgent need to understand the pathobiology of SARS-CoV-2, that might shed light on the causes of long-term symptoms ^1–5^.

RNA viruses are the etiologic agents of many prevalent and lethal human diseases ^6^. It is well documented that despite the completion of their life cycle within the host cell cytoplasm, RNA viruses can induce significant DNA damage and activate the DNA damage response (DDR) pathway. Both events enable viral replication and modulation of host cell functions. Notably, the positive-sense RNA viruses from the family *Coronaviridae* to which the SARS-CoV-2 belongs, and the negative-strand Influenza A virus from the *Orthomyxovirida*e family induces the DDR pathway in host cells ^7–9^. More recently, global phosphorylation mapping ^10^ and ATR DDR inhibitor studies ^5^ indicate that SARS-CoV-2 may also engage the DDR pathway to propagate in host cells.

A classical DNA damage response is mediated by one of the signaling pathways—ATM, ATR, and DNA-PK kinases ^6^. Double strand breaks usually engage the ATM and the DNA-PK pathways, while single-stranded DNA activates the ATR kinase pathway ^11^. These pathways activate specific downstream effectors such as CHK1 by ATR and CHK2 by ATM to trigger a cellular response that allows cells to arrest cell cycle progression to repair damaged DNA. The DDR pathway is a critical component of an intracellular defense mechanism that is activated upon detection of lesions on the DNA to facilitate DNA repair by any of the following: base excision repair (BER), nucleotide excision repair (NER), mismatch repair (MMR), non-homologous end-joining (NHEJ) or homologous recombination (HR) to fix the damaged DNA ^12^. In the event that DNA repair fails, either programmed cell death is induced, or an alternative pathway of DNA damage tolerance or translesion synthesis (TLS) is triggered ^13^ to allow cell survival despite the presence of DNA damage. Additionally, it is known that ATR and ATM-dependent DNA damage responses associate with telomere dysfunction ^14, 15^. Telomeres are specialized DNA-protein structures that play a key role in maintaining genome stability by protecting the ends of chromosomes ^16^. The shelterin protein complex, comprised of six proteins—TRF1, TRF2, POT1, TPP1, TIN2 and Rap1— specifically associate with telomeric sequences to prevent the chromosomal ends from being recognized as double strand breaks ^17^. Depletion or loss of components of the shelterin complex results in telomere shortening ^18^. Literature survey suggests that Epstein-Bar Virus (EBV) infections can also destabilize telomeres by downregulating TRF2 expression ^19^. Whether SARS-CoV-2 might trigger telomere dysfunction is not known. Herein, we investigated the ability of SARS-CoV-2 to impact the DNA damage response and telomere stability in Vero E6 cells.

## Materials and Methods

### Cell Culture and SARS-CoV-2 Infection

SARS-CoV-2 strain 2019-nCoV/USA_USA-WA1/2020 (WA1) was generously provided by Kenneth Plante and the World Reference Center for Emerging Viruses and Arboviruses at the University of Texas Medical Branch and propagated in African green monkey kidney cells (Vero E6) that were kindly provided by J.L. Whitton. All experiments involving infectious SARS-CoV-2 were conducted at the UVM BSL-3 facility under an approved IBC protocol. Vero E6 cells were infected with SARS-CoV-2 WA1 at an MOI of 0.01 and incubated for 48 hours before collecting the cells for further downstream processing.

### RNA Extraction

RNA from infected cell lysates was harvested by incubating infected Vero E6 cells for 10 minutes with RLT buffer (Qiagen) containing 2-Mercaptoethanol (Fisher Scientific). The lysate was mixed thoroughly, followed by addition of one volume of 70% ethanol (Fisher Scientific) and mixed again by pipetting. The lysate was then transferred to the RNeasy spin column provided in the RNeasy kit (Qiagen). The manufacturers protocol was then followed for the rest of the isolation process.

### RT-qPCR and Telomere Length Assay

Isolated RNA was quantified using the Nanodrop-2000 (ThermoFisher), and then diluted using RNase/DNase free water (VWR Life Sciences) until all the concentration of RNA in each sample was 10ng/µL. The iTaq Universal SYBR Green One-Step Kit (Bio-Rad) was used to run the RT-qPCR reactions, using the manufacturer recommended cycling conditions for the StepOnePlus thermal cycler.

The telomere lengths were measured by using the Relative Telomere Length Quantification qPCR kit (ScienCell) which utilizes primers that recognize telomeric repeats (primer Tel) and the resulting gene expression is normalized to the gene expression of a single copy reference (SCR) control that recognizes a 100 bp region on human chromosome 17. 1µL of DNA at a concentration of 1ng/µL was mixed with either 2µL of primer Tel or primer SCR and 10µL PowerTrack SYBR Green Master Mix (ThermoFisher) with the volume of the reaction being brought up to 20µL with ddH_2_O. The cycling conditions for this reaction were 2 minutes at 95°C followed by 40 cycles of 15 seconds at 95°C and 1 minute at 60°C, with the reaction taking place on the StepOnePlus thermal cycler (ThermoFisher), and the average telomere length was calculated using the manufacturer’s instructions. The StepOnePlus software was used to analyze that data and GraphPad Prism 9.0.1 was used for statistical analysis and to the plot the results. All other primers used for RT-qPCR were obtained through ThermoFisher and are listed below.

### Primers

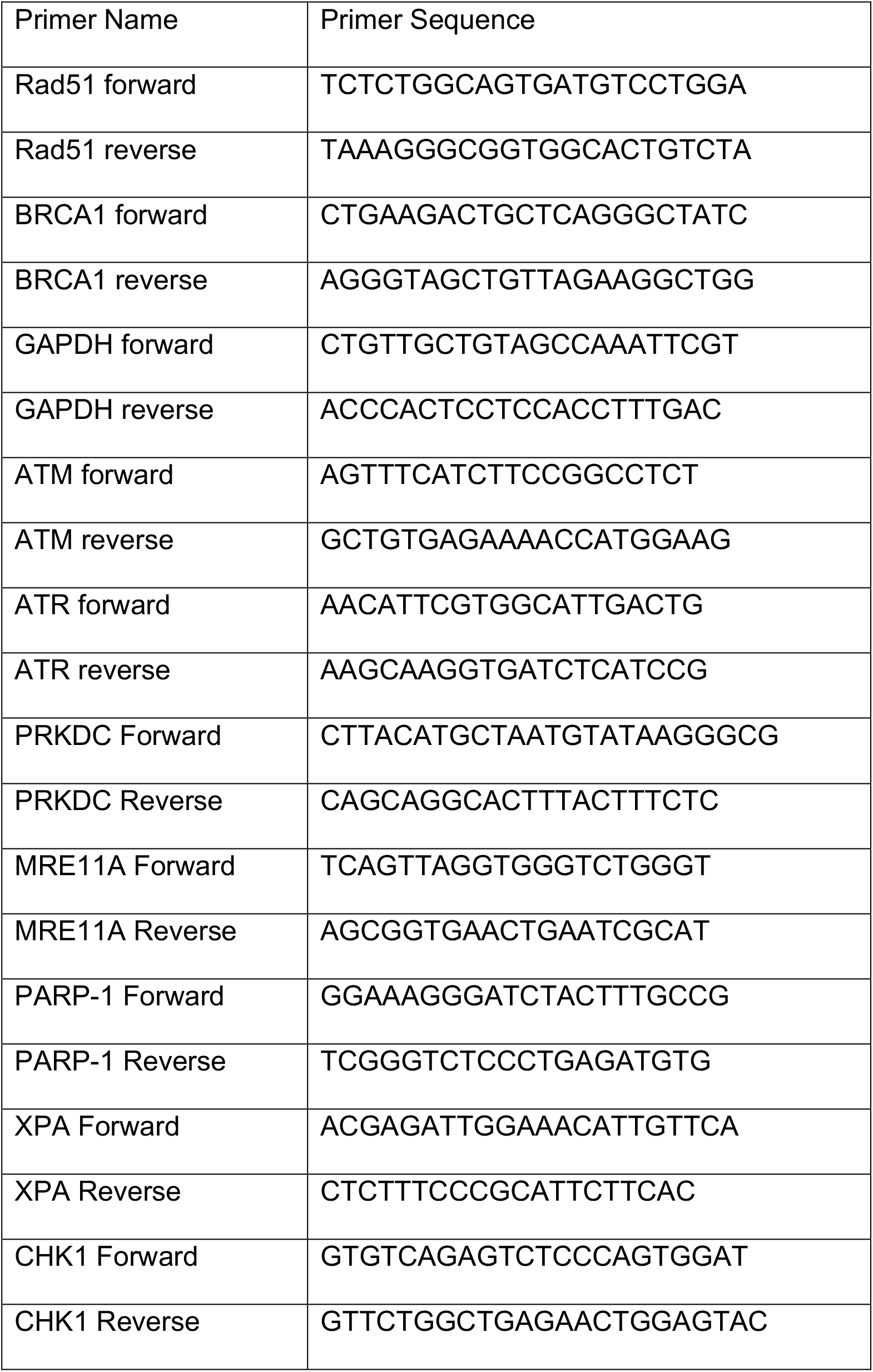

### Immunoblotting and Antibodies

Protein lysates from SARS-CoV-2 infected Vero E6 cells were harvested in 2x Laemmli sample buffer (Bio-Rad) containing 2-Mercaptoethanol (Fisher Scientific). Samples were passed through a 25G syringe five times to reduce viscosity and separated on a 3%-8% NuPAGE Tris-Acetate gel (Thermo Fisher). Proteins were transferred onto a polyvinylidene fluoride membrane (Thermo Scientific) for 90 minutes at 100 volts using the Mini Trans-Blot Electrophoretic Transfer Cell (Bio-Rad). Membranes were blotted with the following antibodies: Actin (Invitrogen, MA1744, Mouse mAb) at 1:1000, CHK1 (Abcam, ab47574, Rabbit pAb) at 1:500, ATR (Abcam, ab2905, Rabbit pAb) at 1:500, phospho-Chk1 (Abcam, ab92630, Rabbit mAb) at 1:500 with 0.01% Tween-20, phospho-ATR (Invitrogen, 720107, Rabbit pAb) at 1:1000, γH2AX (Novus Biological, NB-100-384, Rabbit pAb) at 1:1000, and TRF2 at 1:1000 (Abcam; ab108997). Membranes were washed three times for five minutes each with PBS (Cytia) with 0.1% Tween-20, before and after incubating with secondary antibody for one hour, with donkey anti-mouse (IRDye 680RD, Li-COR BioSciences) or goat anti-rabbit (IRDye 800CW, Li-COR Biosciences), at 1:20,000 with 0.01% SDS and 0.1% Tween-20. Membranes were imaged with the Li-COR Odyssey CLx, and images were analyzed with Image Studio software. Bands indicating our proteins of interest were normalized to Actin, and SARS-CoV-2 treated results were further normalized to the mock controls.

## Results and Discussion

To test the hypothesis that SARS-CoV-2 infection triggers a DDR, we infected Vero E6 cells at an MOI of 0.01 and tested expression of DDR genes after 48 hours (Fig. 1A). At this MOI and exposure time frame, Vero E6 cells produce maximal infectious virus outputs ^20^ and remain infected with SARS-CoV-2, as is indicated the relative N2 transcript levels in the SARS-CoV-2 infected cells (Fig. 1B). Our test panel represented genes that were either previously implicated in host cell response to viral infections such as BRCA1, RAD51, ATM, ATR, PRKDC, MRE11 ^21, 22^, or picked as a logical extension of the key genes in the DNA repair pathways, such as PARP1, XPA, and CHK1. Transcript and western blot analyses showed an activation of the ATR DNA damage response post SARS-CoV-2 infection at 48 hours. We observed a significant increased transcript expression of the ATR and CHK1, the downstream effector molecule of ATR, in addition to the increased phosphorylation of the CHK1 protein, indicative of an activated ATR DNA damage response (Fig. 1D and 2A). Within this context, we did not see an enhanced phosphorylation of the ATR protein or the total ATR protein levels in infected cells (Fig. 2A and 2B), suggesting that the overall increase in ATR levels corresponding to the increased mRNA levels may have occurred prior to our test time of 48 hours. In fact, we observe an overall reduction in both the total ATR and CHK1 protein levels at 48 hours (Fig. 2B). Ascertaining the increased expression of ATR and CHK1 in a time course experiment post SARS-CoV-2 infection will be of interest in future studies. Interestingly, we also observed an increase in H2AX phosphorylation protein, despite a lack of an increase in ATM transcript expression. We conclude that SARS-CoV-2 infection activates the host cell ATR DDR pathway, which could provide an unknown proliferation potential to its infectious cycle ^5^.

**Figure 1.**
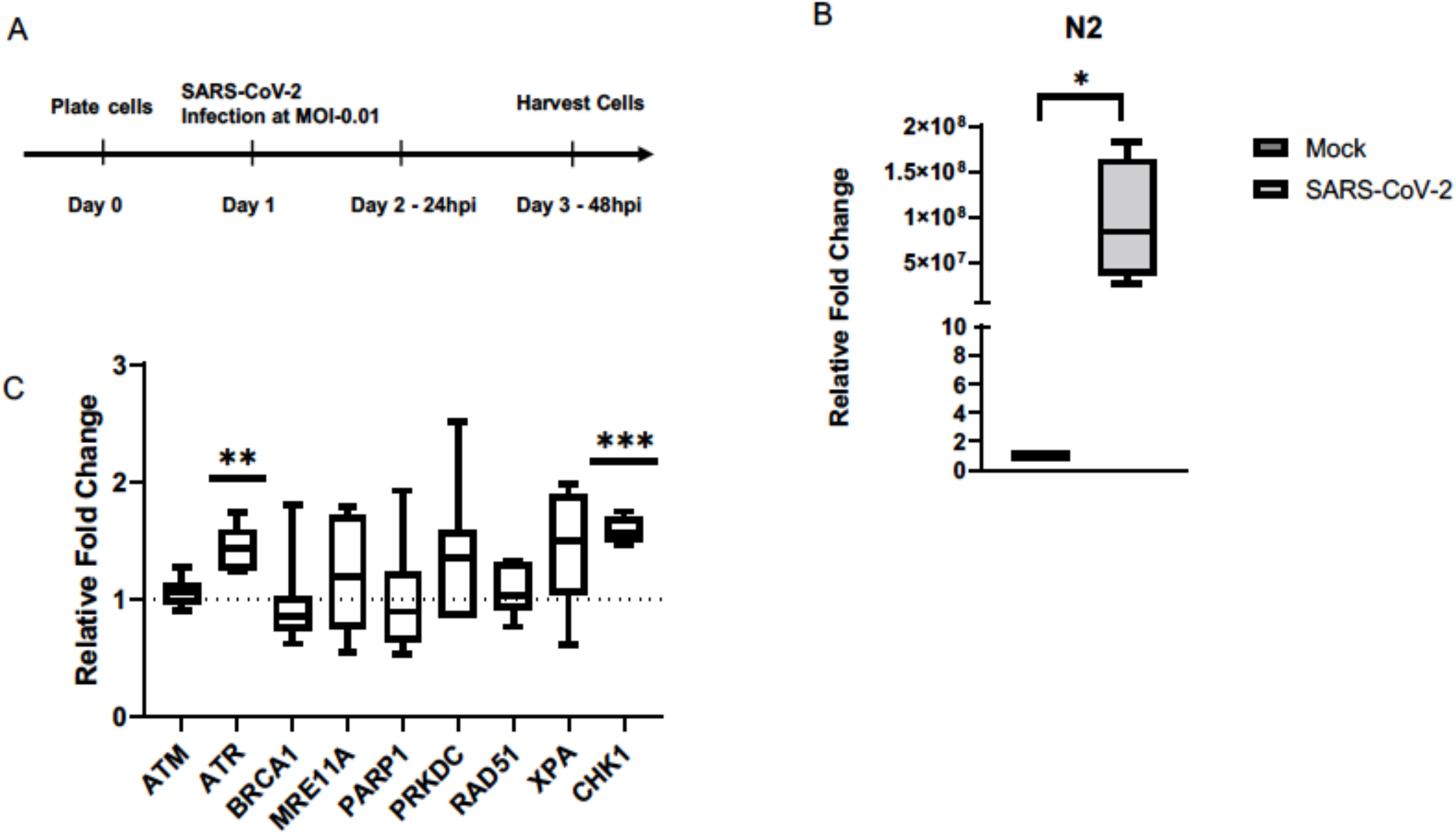
SARS-CoV-2 infection induces expression of ATR and CHK1 in Vero E6 cells. **(A)** Experimental outline of infecting cells with SARS-CoV-2. **(B)** N2 transcript levels were measured and confirmed that there was an active infection in the Vero E6 cells 48 hours after being infected with SARS-CoV-2. **(C)** Relative transcript levels of key DNA damage response genes in Vero E6 cells infected with SARS-CoV-2. (n=3 for N2, n=4 for CHK1, n= 6 for ATM, ATR, and MRE11A; n=7 for PRKDC; n=8 for BRCA1 and PARP1; n=10 for Rad51 and XPA). ^*^P<0.05, ^***^P<0.0005.

**Figure 2.**
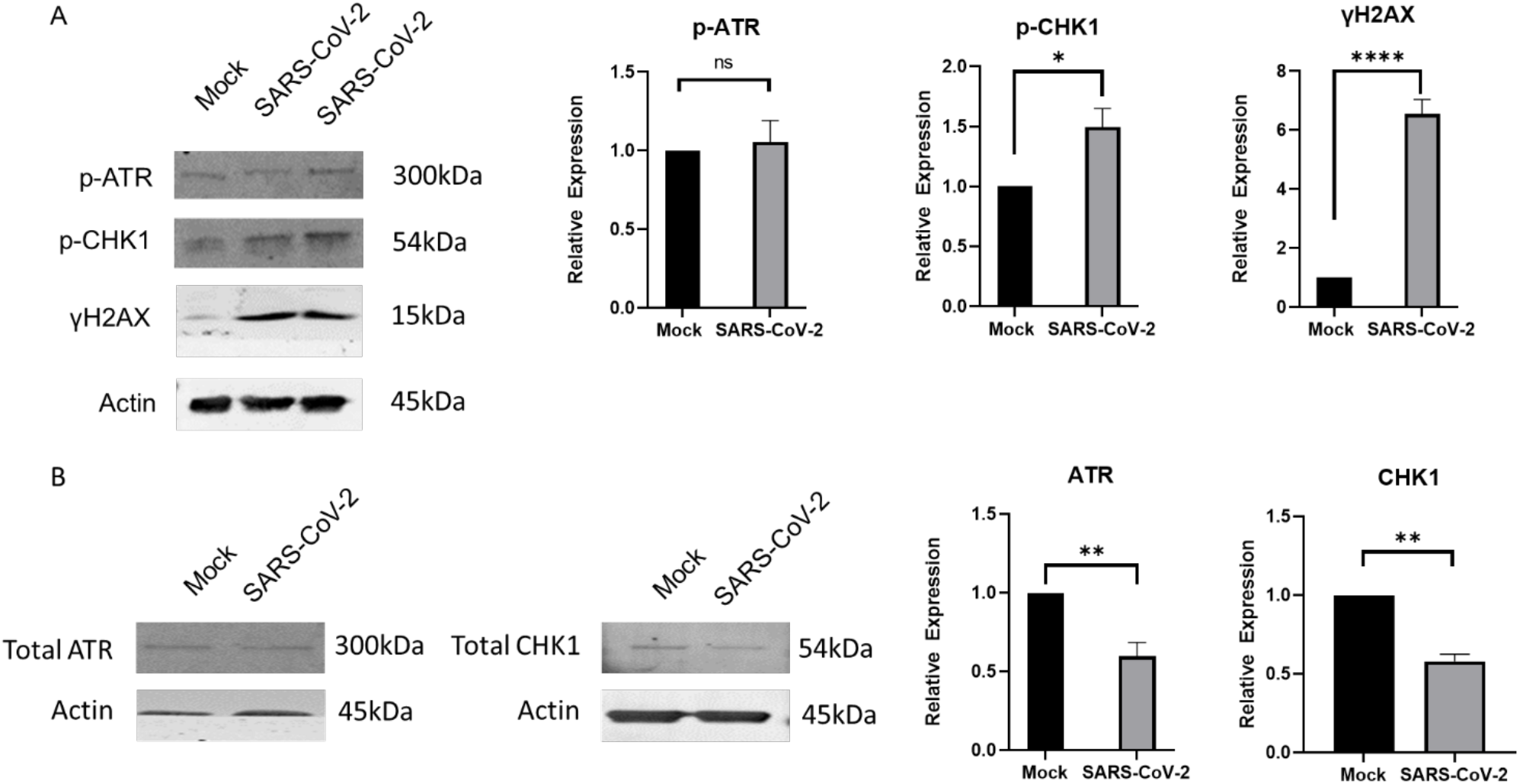
SARS-CoV-2 infection triggers activation of downstream molecules of the ATR DNA damage response. **(A)** Shows representative image of the western blot showing significant increase in phospho-CHK1 and γH2AX expression in SARS-CoV-2 infected cells. Quantification plots showing relative change in expression of p-ATR, p-CHK1, and γH2AX. **(B)** Shows representative images of the western blots showing a significant increase in total CHK1 and ATR levels, as well as quantification plots showing relative change in expression of total CHK1 and ATR. Error bars represent standard error of the mean (n=3 for CHK1 and ATR, n= 10 for γH2AX; n = 12 for p-CHK1; n = 13 for p-ATR). ^*^P<0.05, ^**^P<0.005 and ^****^P<0.000005.

ATR activation due to retroviral infections such as Human immunodeficiency Virus 1 (HIV-1) and Avian Reovirus (ARV) are expectedly driven by creation of double strand breaks during the integration of viral DNA that then leaves behind single strand gaps ^23, 24^. Interestingly, ATR DNA damage response is also a successful strategy to manipulate the cell cycle progression utilized by RNA viruses to propel their infection cycles as reported for another member of the *Coronaviridae* family, infectious bronchitis virus (IBV) ^7, 25^. Further studies are needed to ascertain the utility of ATR DNA damage response in propagation of the SARS-CoV-2 virus in cells.

To quantify possible effects of an activated ATR DNA damage response in infected cells, we measured telomere lengths both in mock and SARS-CoV-2 infected cells. We used the commercially available qPCR-based Relative Telomere Length Quantification kit and compared the relative amplification of the telomere end to the internal control. Strikingly, we observed that SARS-CoV-2 infection shortens the relative length of telomeres compared to mock controls within 48 hours (**Fig. 3A**). In addition, we examined the relative expression of one of the key telomere proteins, TRF2, and found its expression to be significantly suppressed in SARS-CoV-2 infected Vero E6 cells (**Fig. 3B and C**). Whether other cell types, such as rodent or human cells would exhibit a similar telomere phenotype post-SARS-CoV-2 infection needs to be ascertained. TRF2 is one of the most important shelterin complex proteins that ensure telomere end protection from exonuclease degradation that maintains proper telomere length and genome integrity ^26^. Depletion of TRF2 results in telomere fusions and, or, shortening depending upon the physiological context ^27^. A literature survey revealed that EBV (a DNA virus that causes mononucleosis) infections can destabilize telomeres via downregulation of TRF2 ^19^. It is not known how the RNA viruses, particularly SARS-CoV-2, could modulate TRF2 expression and destabilize telomere lengths. Short telomeres associate with the onset of senescence, and a host of other debilitating conditions ^28^.

**Figure 3.**
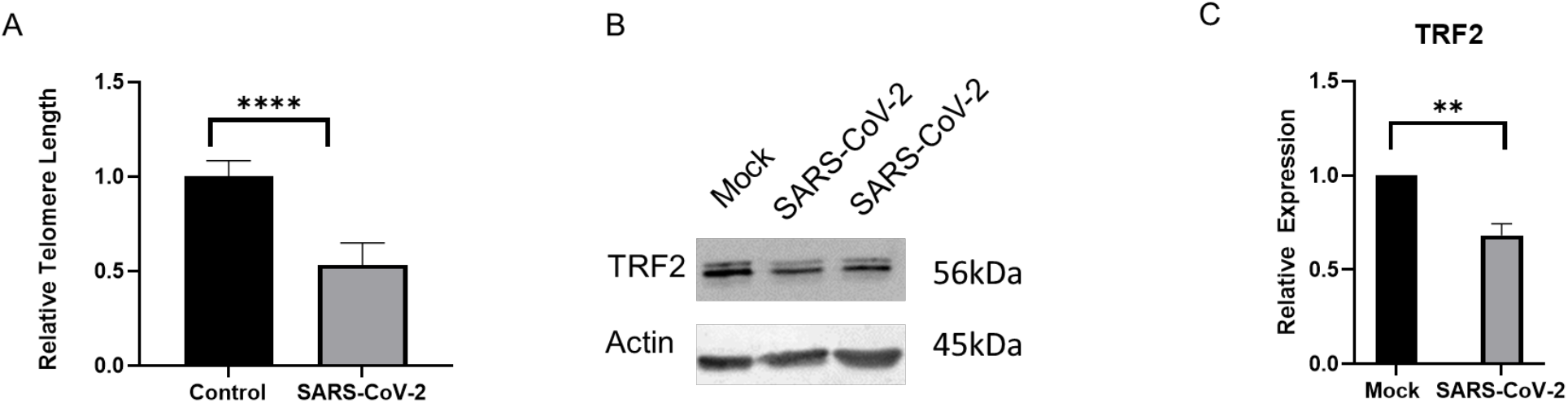
SARS-CoV-2 infection causes telomere dysfunction in Vero E6 cells. **(A)** Shows relative differences in telomere lengths in SARS-CoV-2 infected cells versus the controls. The Relative Telomere Length Quantification qPCR kit from ScienCell was used for this experiment. **(B)** Shows a representative image of the western blot of the TRF2 protein in mock and SARS-CoV-2 infected lysates. **(C)** Shows quantification of TRF2 expression from B. Error bars represent standard error of the mean (n= 10 for TRF2 and n=14 for Telomeres). ^****^P<0.000005 and ^**^P<0.005.

SARS-CoV-2 has made a lasting impact across the globe infecting millions of people, and spurred concern due to long-term health consequences. With the advent of the continuously evolving strains, studying the pathobiological consequences of this infection in recovered patients is vital. This study suggests that in Vero E6 cells, SARS-CoV-2 infection triggers an ATR DNA damage response and affects telomere function. Both ATR activation and telomere instability are associated with genome instability. Further studies are required to expand these findings and ascertain the clinical implications of these results.

## Author Contributions

JAV conducted qPCRs and telomere studies with help from AM. JD conducted western blots with help of ENL. AW carried-out SARS-CoV2 infections with guidance from JWB. PZ helped with data analysis and provided critical intellectual input. NC conceived the project, designed experiments, and wrote the manuscript with help from all authors.

## Acknowledgments

This work was supported in part by the University of Vermont Cancer Center summer fellowship to JAV, DUSRA research award to ENL, the University of Vermont Office of the Vice President for Research to JWB, University of Vermont Cancer Center Pilot project grant to NC, the NIH T32 AI055402 training grant to AW, the COBRE Pilot project grant from the Vermont Institute of Immunology and infectious diseases to NC, and University of Vermont Office of the Vice President for Research—Bellinger research funds to NC.

